# PopPAnTe: population and pedigree association testing for quantitative data

**DOI:** 10.1101/084871

**Authors:** Alessia Visconti, Mashael Al-Shafai, Wadha A. Al Muftah, Shaza B. Zaghlool, Massimo Mangino, Karsten Suhre, Mario Falchi

## Abstract

Family-based designs, from twin studies to isolated populations with their complex genealogical data, are a valuable resource for genetic studies of heritable molecular biomarkers. Existing software for family-based studies have mainly focused on facilitating association between response phenotypes and genetic markers, and no user-friendly tools are at present available to straightforwardly extend association studies in related samples to large datasets of generic quantitative data, as those generated by current -*omics* technologies.

We developed PopPAnTe, a user-friendly Java program, which evaluates the association of quantitative data in related samples. Additionally, PopPAnTe implements data pre and post processing, region based testing, and empirical assessment of associations.

PopPAnTe is an integrated and flexible framework for pairwise association testing in related samples with a large number of predictors and response variables. It works either with family data of any size and complexity, or, when the genealogical information is unknown, it uses genetic similarity information between individuals as those inferred from genome-wide genetic data. It can therefore be particularly useful in facilitating usage of biobank data collections from population isolates when extensive genealogical information is missing.

## 1 Introduction

Family-based designs, from complex genealogical structure to twin studies, are a valuable resource for genetic studies. The primary aim of currently-available software accounting for population substructure and/or relatedness in the statistical model (*e.g.*, EMMA [1], Merlin [2], GenABEL [3], QTDT [4]) is to evaluate the association between genetic SNP markers and response phenotypes and, to date, very few tools are available to test the association of large quantitative datasets generated by high-throughput -*omics* technologies (*e.g.*, epigenomic versus metabolomic data, or transcriptomic versus metagenomic data) in familial samples. For instance, although modelling of genealogical data can be performed by the *coxme* [5] and *kinship2* [6] R packages, R is not a particularly efficient environment to carry out hundred of thousands or millions tests.

We have implemented a user-friendly Java program, PopPAnTe, to perform exact association tests between large quantitative datasets in family-based studies. Relationships between individuals can be described either by known pedigrees of any size and complexity or by genetic similarity matrices (GSMs) inferred from genome-wide genetic data [7]. Pedigree-based and pedigree-free relatedness can show some discordance, especially when some degree of hidden relatedness or population substructure is observed in the data and extensive genealogical information is missing or incomplete. For instance, genealogical information going back more than three or four generations may be difficult to be retrieved for individuals recruited in large-scale biobank started in genetic isolates such as those from the Middle East.

## Implementation

PopPAnTe assesses the relationship between quantitative dependent variables (*responses*) and quantitative independent variables (*predictors*) within a variance components framework in order to model the resemblance among relatives.

The association of a single predictor with a single response variable is described as

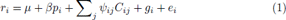

where *r_i_* represents the response value for the *i*-th individual, *μ* the response mean, *β* the estimate of the predictor value *p_i_*, 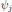 the estimate of the *j*-th covariate *C*, and *g_i_* and *e_i_* the polygenic and environmental effect, respectively.

The total response variance is partitioned into polygenic and environmental variances (the latter including also measurement errors), and the variance-covariance matrix is calculated as

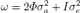

where *ɸ* is the relatedness matrix between each pair of individuals, *I* is the identity matrix, and *σ*_*a*_^2^ and *σ*_*e*_^2^ are the additive genetic and environmental variance, respectively.

Within the same framework, PopPAnTe allows the evaluation of the narrow heritability of any quantitative response variable included in the analysis.

The significance of the association is calculated using a formal likelihood-ratio test comparing the likelihood of the alternative model described in equation (1) to the likelihood of a null model where the effect of the predictor is constrained to zero.

PopPAnTe implements an exact linear mixed model equivalent to that implemented in the QTDT software [4].

To speed-up the evaluation, PopPAnTe clusters variables having the same pattern of missingness (*i.e.*, the same missing values in a subset of individuals), then evaluates the likelihood of the null model once, and reuses the value to assess every variable included in the same cluster. PopPAnTe also allows the evaluation of empirical p-values by randomly permuting the predictor values among subjects and re-assessing the association under the null hypothesis. When genealogical information is provided as input, predictor values are randomly permuted within families in order to preserve the phenotypic correlation between family members. To speed up performance, PopPAnTe implements an adaptive permutation approach [8], stopping the generation of randomly permuted samples earlier when there is little or no evidence of significance.

### Pedigree versus Population analysis

When genealogical information is available, PopPAnTe evaluates the relatedness matrix from the known pedigree relationships using a recursive procedure and assuming pedigree founders as unrelated [9]. This results in a variance-covariance matrix that is usually both symmetric and semi-positive definite. Therefore, the maximum likelihood estimates of the variance components can be assessed through efficient Cholesky decomposition.

When the genealogical information is not available, a GSM can be estimated from genome-wide genetic data with any of several well-established tools, such as PLINK [10], GCTA [11], or LDAK [12], and given as input to PopPAnTe. The property of positive-definiteness does not always hold for GSMs. A bending procedure [13] is used by default to transform the matrix when it is not positive semi-definite –but the user has the option to use a LU decomposition instead. Additionally, PopPAnTe implements the QR decomposition to solve the rare cases where the variance-covariance matrix is not invertible and neither the Cholesky nor the LU decompositions can be used.

To speed-up the evaluation of the variance components, PopPAnTe allows the user to set an arbitrary threshold below which individuals can be considered as unrelated. Otherwise, the user has the options of using the value of expected kinship between second or third cousins [14,15].

### Region-based testing

When predictors can be ordered in space, as it is for instance for gene expression or epigenetic markers, PopPAnTe allows the computation of region-based association tests by gathering information from flanking predictors included in a sliding window of user-defined size, whose values are replaced by their first principal component. By definition the first principal component accounts for as much of the variability in the data as possible, and can thus be used to summarise the joint distribution of all variables included in a given region for gene- or region-based association studies (*e.g.*, [16,17]).

### Data pre-and post-processing

Quantile normalisation [18] can be automatically applied to improve normality of both response variables and predictors. Moreover, PopPAnTe implements two approaches to correct the association test for unwanted biological and technical variability (*e.g.*, batch effects). When the source of the confounders is known, it can be directly included in the association model. To deal with unknown sources of biological and technical co-variation, PopPAnTe can integrate into the association model the principal components that are required to explain a user-specified percentage of variation.

PopPAnTe implements the Benjamini-Hochberg procedure (BH step-up procedure) to control the false discovery rate [19], and, to aid in results interpretation and further analyses, it generates basic Quantile-Quantile and Manhattan plots – the latter only when genomic data that can be ordered in space (*e.g.*, CpG loci) are used as predictors.

Finally, when the genealogical information is available, to determine whether an association has been generated by a uniform contribution of all the families within the sample, or by a strong contribution of a small number of families, PopPAnTe reports, for each test, the percentage of families showing a positive contribution and the Gini coefficient [20] assessed on family contribution to the χ^2^ statistics.

## Results and Discussion

We carried out a simulation study to estimate PopPAnTe's computational requirements. Moreover, we also present two real-world case studies (an outbred and an inbred sample), showing the results obtained through both pedigree-based kinship matrix and GSM (evaluated using different software).

### Simulation study

We simulated quantitative response and predictor variables in three-generation families (maternal and paternal grandparents, parents, and two offspring).

In the first simulated scenario, we aimed to test the relationship between running time and number of subjects included in the analysis. Therefore, we simulated 11 independent datasets including an increasing number of three-generation families (from 10 to 1,000, thus comprising 80 to 8,000 individuals) and one response and one predictor variable for all simulated subjects.

In the second simulated scenario, we aimed to test the relationship between running time and number of variables included in the analysis. Consequently, we fixed the number of families included in each dataset (125 families, corresponding to 1,000 individuals) and generated 7 independent datasets with one response and an increasing number of predictors ranging from one to 10,000.

Both scenarios were simulated 100 times and the median time necessary for the testing step recorded. Simulations were performed on a Mac BookPro 2.3GHz, Intel Core i7, 16GB RAM; Java version 1.7. Default parameters were used for the Java Virtual Machine (1GB of memory, 1 thread).

Tables 1 and 2 show the median running time for each simulated scenario. We observed a linear relationship between running time and both samples size and number of tests. As expected, when multiple tests were performed (second scenario), the per-test running time decreased, due to the fact that PopPAnTe clusters variables having the same pattern of missingness and evaluates the likelihood of the null model only once.

**Table 1.**
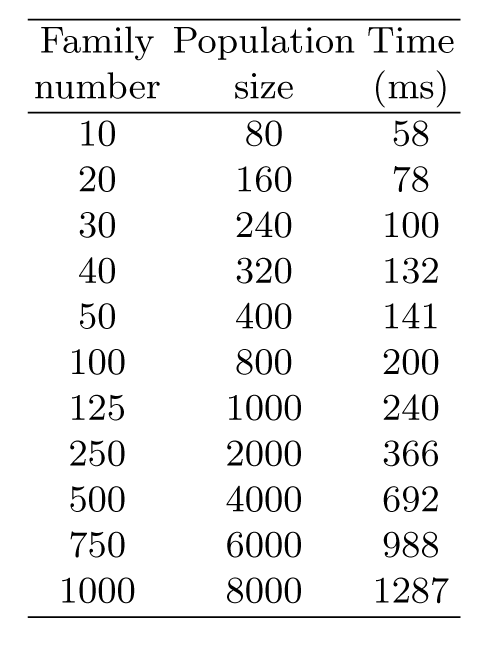
Results of the first simulated scenario. One response and one predictor variable were simulated for each subject. Each dataset was simulated 100 times and the median time necessary for the testing step reported.

**Table 2.** Results of the second simulated scenario. The number of families included in each dataset was fixed (125 families, corresponding to 1,000 individuals). Each dataset was simulated 100 times and the median time necessary for the testing step reported.

| Number of Predictors | Time (s) |
| --- | --- |
| 1 | 0.24 |
| 10 | 0.94 |
| 100 | 7.74 |
| 500 | 37.67 |
| 1000 | 47.00 |
| 2500 | 119.00 |
| 5000 | 238.50 |
| 10000 | 479.00 |

### Case study 1: Epigenome-wide Association Study in a Qatari Family Study

We carried out an epigenome-wide association study of body mass index (BMI) using extended families from Qatar. The Qatari population is an isolated inbred population characterised by a large number of consanguineous families [21]. A detailed description of the subjects and methylation data included in this study has been previously reported in Zaghlool *et al.* [22]. Briefly, we used genome-wide methylation and SNP data generated from whole blood on the Infinium HumanMethylation450 Bead-Chip (Illumina Inc, San Diego, CA) and the Illumina HumanOmni2.5-8M BeadChip, respectively. DNA methylation Beta-values were measured for 123 individuals, 88 with both genotype and BMI data in 13 multigenerational families. We used GCTA to calculate a GSM between pairs of individuals using all autosomal SNP markers with minor allele frequency > 0:01. We compared heritability estimates of the methylation values at CpG loci and their association with BMI in the Qatari family study using either the family information or the inferred GSM. Age, sex, and cell-type proportions as estimated using the Houseman method [23] were included in the model as fixed effects. We observed a very high concordance correlation coefficient [24] of the effect size estimates for the association between CpG methylation states and BMI (*rβ* = 0:99; Figure 1, left), as well as of the CpG-speci c component of genetic and environmental variances (*r*_*a*_^2^, = 0:99 and *r_e_*^2^ = 0:90, respectively; Figure 1, right).

**Fig. 1.**
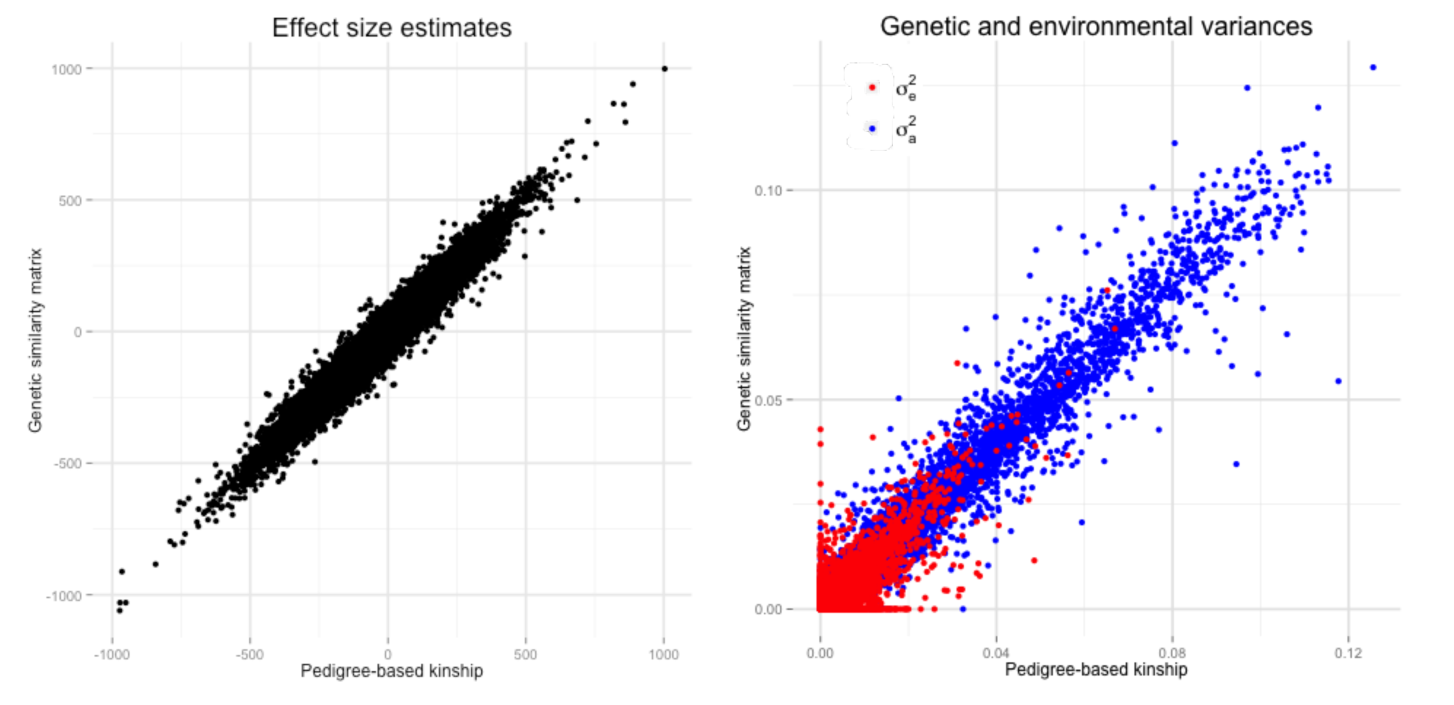
Epigenome-wide Association Study in a Qatari Family Study. Comparison of the results obtained in the Epigenome-wide association studies when the relatedness between subjects was evaluated using the family structures and when it was inferred from genome-wide SNPs by means of GCTA [11]. Left panel: effect size estimates for the association between CpG methylation status and BMI. Right panel: estimated genetic ( *σ*_*a*_^2^, in blue) and environmental ( *σ*_*e*_^2^, in red) variances.

### Case study 2: Transcriptome-wide Association Study in UK Twins

In the second case study, we carried out a transcriptome-wide association study with BMI in a cohort of healthy female Caucasians twins. The TwinsUK adult twin registry includes about 12,000 subjects, predominately females [25]. Genotyping of the TwinsUK dataset was performed with a combination of Illumina HumanHap300, HumanHap610Q, 1M-Duo and 1.2MDuo 1M chips and imputation was performed using the IMPUTE software package (v2), as previously described [26]. Expression profiling in subcutaneous adipose tissue was measured using Illumina Human HT-12 V3 BeadChips for 825 female individuals [27], 778 of whom had both genotype data and BMI information. We used LDAK to calculate a GSM based on allelic correlation across autosomes. Before calculation, we excluded SNPs with minor allele frequency < 0.01. We compared effect size estimates of the gene expression profiles versus BMI, using either the family information or the inferred GSM. Age was included in the model as a fixed effect. We observed a very high concordance correlation coefficient of the effect size estimates for the association between gene expression levels and BMI (*rβ* = 0:99; Figure 2, left)), as well as of the genespecific genetic component of genetic and environmental variances (*r_a_*^2^ = 0:99 and *r_e_*^2^ = 0:99, respectively; Figure 2, right).

**Fig. 2.**
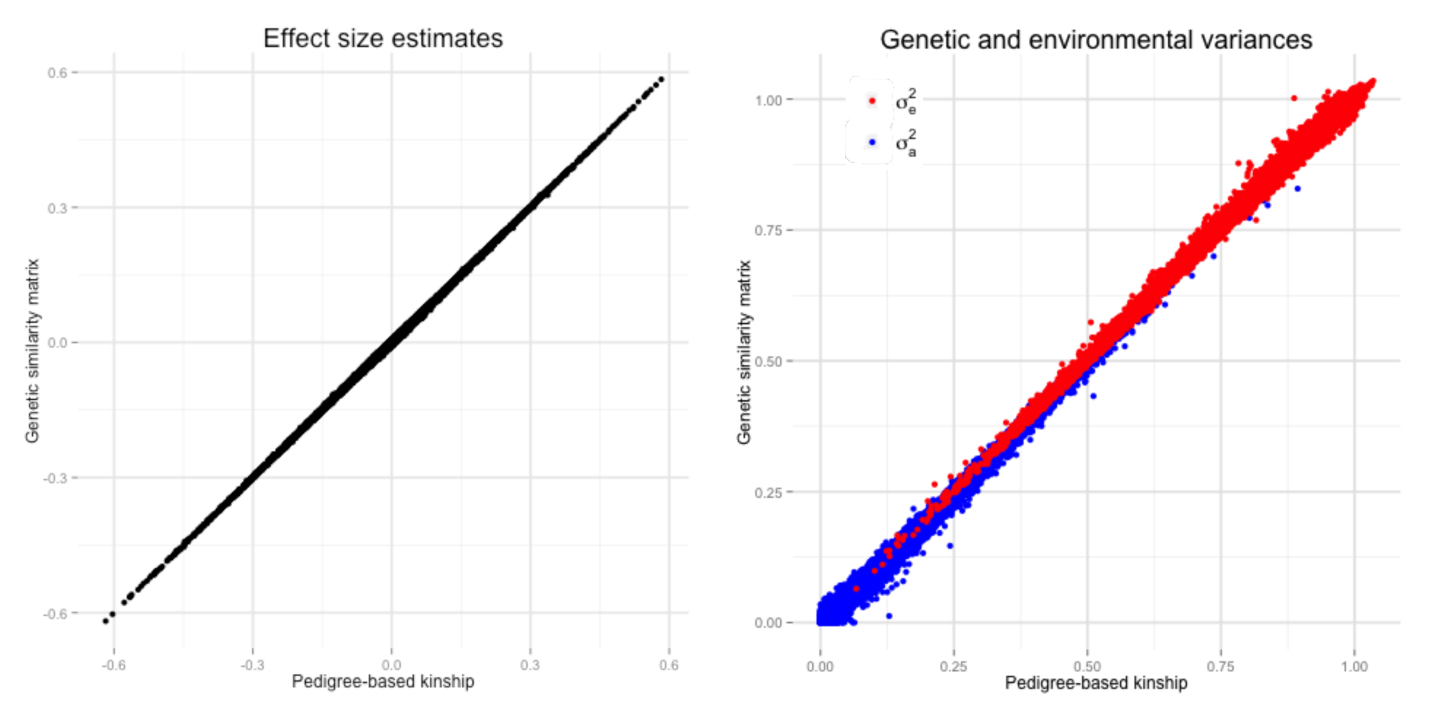
Trascriptome-wide Association Study in UK Twins. Comparison of the results obtained in the transcriptome-wide association when the relatedness between subjects was evaluated using the family structures and when it was inferred from genome-wide SNPs by means of LDAK [12]. Left panel: effect size estimates for the association between gene expression levels and BMI. Right panel: estimated genetic ( *σ*_*a*_^2^, in blue) and environmental (*σ*_*e*_^2^, in red) variances.

## 2 Conclusions

PopPAnTe is a user-friendly platform-independent Java program that enables pairwise association testing of large numbers of predictor and response variables in related samples. PopPAnTe uses either known pedigree structures or GSMs inferred from genome-wide genetic data, allowing the user to select the best approach according to the available data. When genome-wide genetic data are available, it may be advisable to use the GSM instead of the expected kinship matrix calculated using the genealogical information [28,29]. PopPAnTe can thus also facilitate the usage of biobank collections from population isolates when extensive genealogical information is missing.

## Author's contributions

AV, KS, and MF designed the software. AV implemented PopPAnTe, and performed the computational experiments. MAS, WAAM, SBZ, and KS generated the Qatari family study data. MM provided the data for the TwinsUK cohort, and tested the software. AV and MF wrote the manuscript. All authors read and approved the final manuscript.

## Acknowledgements

We thank the Qatar Diabetes Association (QDA) and Cindy McKeon for their help in sample recruitment. AV and MF thank S. Burbidge and M. Harvey of the Imperial College High-Performance Computing service for their assistance, Doug Speed for his valuable suggestions, and Julia El-Sayed Moustafa for her constructive comments on the manuscript. We are grateful to all study participants for their contribution to this research study. The two studies were conducted in concordance with the Helsinki declaration of ethical principles for medical research involving human subjects. The studies were approved by the relevant institutional review boards in Qatar (Institutional Review Board of Weill Cornell Medical College in Qatar, ethical approval numbers 2012003 and 20120025), and in the UK (Guys and St. Thomas Hospital Ethics Committee). Written informed consent was obtained from every participant in each study.

### Funding

This work was supported by the Qatar Science Leadership Program at the Research Division, a program funded by the Qatar Foundation, and by Bioinformatics and Virtual Metabolomics Core of Weill Cornell Medical College in Qatar. MF and AV are supported by the British Skin Foundation grant 5044i. MF is supported by MRC grant MR/K01353X/1. The TwinsUK study was funded by the Wellcome Trust; European Communitys Seventh Framework Programme (FP7/2007-2013). The study also receives support from the National Institute for Health Research (NIHR)- funded BioResource, Clinical Research Facility and Biomedical Research Centre based at Guys and St Thomas NHS Foundation Trust in partnership with Kings College London. SNP Genotyping was performed by The Wellcome Trust Sanger Institute and National Eye Institute via NIH/CIDR.

## Availability and requirements

**Project name:** PopPAnTe

**Project home page:** https://sites.google.com/site/populationgenomics/poppante

**Operating system(s):** Platform independent

**Programming language:** Java

**Other requirements:** Java 1.7 or higher

**License:** GNU GPL 3 or higher

**Any restrictions to use by non-academics:** None

